# Human Satellite 1 (HSAT1) analysis provides novel evidence of pericentromeric transcription

**DOI:** 10.1101/2022.08.11.503625

**Authors:** Mariana Lopes, Sandra Louzada, Daniela Ferreira, Gabriela Veríssimo, Daniel Eleutério, Margarida Gama-Carvalho, Raquel Chaves

## Abstract

Pericentromeric regions of human chromosomes are composed of tandem-repeated and highly organized sequences named satellite DNAs. Although being known for a long time as the most AT-rich fraction of the human genome, classical satellite HSAT1 has been disregarded in genomic and transcriptional studies, falling behind other human satellites in terms of knowledge. The path followed herein trails with HSAT1 isolation and cloning, followed by *in silico* analysis. Monomer copy number and expression data was obtained in a wide variety of human cell lines, with greatly varying profiles in tumoral/non-tumoral samples. HSAT1 was mapped in human chromosomes and applied in *in situ* transcriptional assays. Additionally, it was possible to observe the nuclear organization of HSAT1 transcripts and further characterize them by 3’ RACE-Seq. Size-varying polyadenylated HSAT1 transcripts were detected, which possibly accounts for the intricate regulation of alternative polyadenylation. As far as we know, this work pioneers HSAT1 transcription studies. With the emergence of new human genome assemblies, acrocentric pericentromeres are becoming relevant characters in disease and other biological contexts. HSAT1 sequences and associated noncoding RNAs will most certainly prove significant in the future of HSAT research.

## Introduction

Satellite DNA (satDNA) sequences consistently organize in arrays of tandem repeats, preferentially located at (peri)centromeric and subtelomeric heterochromatin (Yunis and Yasmineh 1971; Warburton et al. 2008). Historically, these sequences were acknowledged by distinguishable satellite bands in cesium chloride gradients of human genomic DNA (Kit 1961) and termed classical satellites I, II, and III (Choo 1997; Lee et al. 1997), today known as human satellite families HSAT1, HSAT2, and HSAT3. The HSAT1 (also known as SATI) family is composed of two alternating repeat unit types, A (17 bp) and B (25 bp), combined in a 42 bp monomer. This satellite was first described with a probe (pTRI-6) locating at chromosomes 3 and 4, and acrocentric chromosomes (chr13, 14, 15, 21, and 22) (Kalitsis et al. 1993; Trowell et al. 1993).

The human reference genome (current patch GRCh38.p14) is still an undeniable hostage of assembly issues related with acrocentric HOR/pericentromeric sequence sharing, evidently under- or misrepresenting satDNA sequences (Lopes et al. 2021). From a closer look into Dfam (Hubley et al. 2016) (https://dfam.org) or Repbase (Bao et al. 2015) (https://www.girinst.org/repbase/), HSAT1 can be found in two different annotation types: SAR (DF0001062.4), firstly acknowledged to repeat in 42 bp units (Prosser et al. 1986), annotated in chromosomes 4, 8, 14, 15, and 22 (hg38); and HSATI (DF0000210.4), formerly identified as male-specific and containing one Alu family member (Frommer et al. 1984). Addressing the high number of gaps in the human reference assembly related with acrocentric p-arms, the Telomere-to-Telomere (T2T) consortium has recently released the T2T-CHM13 human genome assembly (Nurk et al. 2022), of which HSAT1 constitutes 0,47% of the total sequence (Altemose et al. 2022). In this work, HSAT1 was re-classified into HSAT1A and HSAT1B elements. HSAT1A (corresponding to SAR), the main scope of the present paper, was predominantly found to form longer, 378 bp tandem 9-mer repeats of the 42bp monomers (Altemose et al. 2022).

Despite being a part of constitutive heterochromatin, Human Satellites (HSATs) are not transcriptionally inactive (Ugarkovic 2005). HSAT transcription into satellite noncoding RNAs (satncRNAs) is reported as bidirectional and promoted by RNA polymerase II (Pol II) (Bury et al. 2020). Pericentromeric satellite transcription was reported as strand-specific depending on cell state (e.g., stress *versus* senescence) (Jolly et al. 2004; Enukashvily et al. 2007). As products of Pol II transcription, ncRNAs regulation is dependent of cotranscriptionally-occurring RNA processing. Polyadenylation consists in the 3’ processing of mRNAs or ncRNAs through the addition of poly(A) tails, known to influence RNA stability and transport (Lewis et al. 1995; Wickens et al. 1997). Shortly, when poly(A) signals emerge during nascent transcription, 3’-end cleavage and polyadenylation (CPA) complex is recruited, inducing Pol II slowdown and transcription termination. Transcripts of variable lengths can therefore result from alternative polyadenylation (APA) using different poly(A) sites (PASs) (Proudfoot 2016). Premature CPA (PCPA) can result from the selection of proximal PASs, especially in highly proliferative cells (Tian and Manley 2013). PCPA is strictly balanced by its suppressing counterpart - a process termed telescripting, which essentially assures full transcript length. Therefore, both mRNA and ncRNA transcript variability depends on the regulation of the PCPA-telescripting duality (Di et al. 2019).

Many ncRNAs are transcriptionally dysregulated in disease states: in transcript levels, (Deveson et al. 2017), and potentially in transcript processing, or both. Increased satDNA expression has been related with genomic instability (Chatterjee and Sengupta 2021). Whilst preserving chromosome integrity (Plohl et al. 2008; Bierhoff et al. 2014; Ershova et al. 2019), tandem repeats can potentially represent beacons of instability in cancer genomes (de Lima et al. 2021). Chromosome breaks tend to occur in pericentromeric satellite regions (Black and Giunta 2018), possibly altering the regulation of transcription of satellite sequences (Ferreira et al. 2015; Puppo et al. 2020). By its turn, available pericentromeric chromatin in cancer might also predispose chromosomes to break (Louzada et al. 2020). Hence, cancer progression can be related with the emergence of chromosomal alterations, caused by (or causing) changes in genomic architecture and transcription deregulation in noncoding regions (Spielmann et al. 2018; Louzada et al. 2020).

NcRNAs, and more precisely satncRNAs, expression is underrepresented in transcriptomic data (Bu et al. 2015), as ncRNAs seem to pose methodological and analytic challenges, relating with their complex diversity and repetitive nature (Cao et al. 2018). Thus, the function of the human noncoding genome and satellite transcription has been gradually addressed. Centromeric αSAT (Alpha Satellite) transcripts in particular have been considered vital for kinetochore stabilization and centromere cohesion (Liu et al. 2015; McNulty et al. 2017; Smurova and De Wulf 2018; Chen et al. 2021). The progressive description of pericentromeric HSAT2 and HSAT3 transcripts has elevated their status to essential in several cellular contexts (Eymery et al. 2009a), like the formation and regulation of heterochromatin (Saksouk et al. 2015; Johnson et al. 2017), aging (Enukashvily et al. 2007; Mendez-Bermudez et al. 2021), response to stress (Jolly et al. 2004; Goenka et al. 2016; Ninomiya et al. 2020), differentiation (Yandim and Karakülah 2019; Dobrynin et al. 2020), and cancer (Hall et al. 2017; Nogalski and Shenk 2020; Chatterjee and Sengupta 2021).

Comprehensive work describing pericentromeric HSAT2 and HSAT3 has been presented over recent years. However, HSAT1 lacks molecular and cytogenomic studies. To characterize HSAT1, we present copy number variation and transcriptional analysis in distinct cell lines (tumoral and non-tumoral), along with single-cell analysis by RNA-Fluorescence *in situ* hybridization (RNA-FISH). This study is coupled with cytogenetic mapping and immunofluorescence, as well as *in silico* HSAT1 assessment. We also performed 3’ RACE-seq and pointed some hints to the inclusion of HSAT1 into the ncRNA landscape. To the best of our knowledge, this work constitutes, so far, the deepest analysis of pericentromeric HSAT1, the most AT-rich fraction of the human genome.

## Results

### HSAT1 isolation, copy number analysis, and expression profiling

The characterization of HSAT1 is hampered by the low amount of information related with this satellite and its sequence, stemming in part from technical constrains associated to its AT-richness and low read coverage in sequencing studies. Therefore, we decided to take a more classic approach and begin by performing HSAT1 PCR isolation followed by cloning and Sanger sequencing (Supplemental Table S1). The 83 obtained clones (Accession number: OP172545 - OP172627) presented an overall high degree of sequence similarity, with a mean of 87,9% identity. Sequence analysis with the Tandem Repeats Finder software (Benson 1999), revealed a 42 bp repeat pattern (or multiples of) (Figure 1A), and an average GC content of 23%, characteristic of HSAT1 AT-rich sequences.

**Figure 1.**
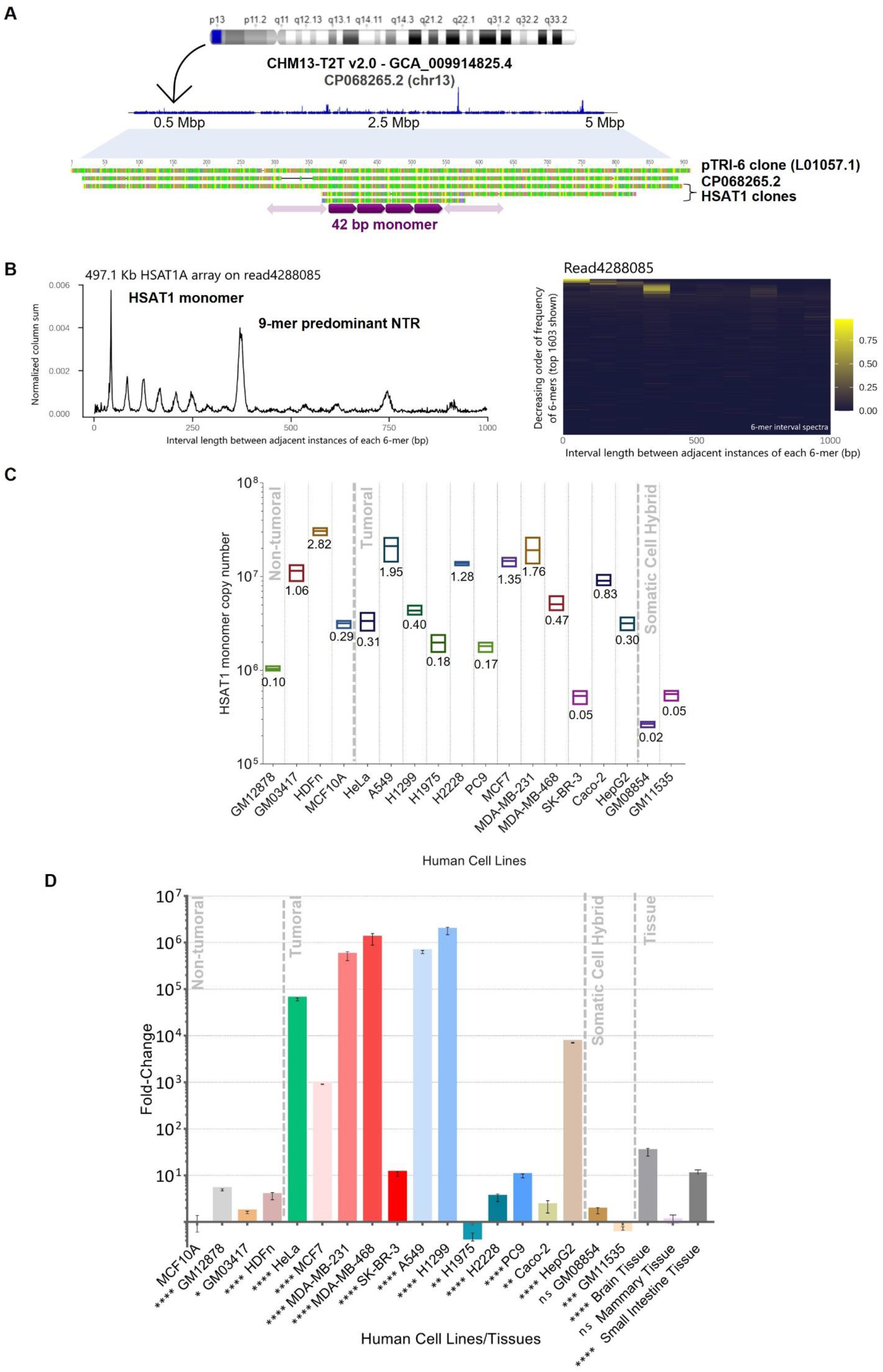
HSAT1 sequence analysis and copy number/expression evaluation. **(A)** – Obtained HSAT1 clones (3-5) were analyzed in Tandem Repeats Finder and proved to be systematically composed of 42 bp repeats (6). HSAT1 clone was BLAST searched against CHM13-T2T v2.0 (GenBank assembly accession GCA_009914755.4) and filtrated hits were mapped into chromosomes. HSAT1 BLAST hits are represented (in blue) in CHM13-T2T chromosome 13 (CP068265.2), reported to have a large HSAT1A array (Altemose et al. 2022). The ideogram was adapted from Ensembl genome browser. *In silico* mapping of HSAT1 hits was performed in Geneious. Concatenated hits are observable in a 5 Mb extent. HSAT1 clones, HSA13 T2T HSAT1A array (1) (Lee et al. 2022; Nurk et al. 2022), and pTRI-6 (2) (L01057.1) sequence stretches were aligned (Geneious alignment, default parameters). **(B)** – HSAT1 periodicity spectrum and heatmap in GM12878 sequencing data. NTRprism reveals two predominant peaks: one corresponding to HSAT1 monomer and the second to a 9-mer higher repeat. **(C1)** – HSAT1 monomer copy number absolute quantification in several human cell lines. Values are mean ± SD. Statistical analysis is detailed in Supplemental Figure S2. HSAT1 estimation in % of the human haploid genome (*) (bp/ total bp). **(D)** – HSAT1 ncRNA relative quantification by RT-qPCR in fold change (MCF10A set as reference). Values are mean ± SD. *P≤0.05, **P≤0.01, ***P≤0.001, ****P≤0.0001, ns - not statistically significant (one-way ANOVA with Tukey ‘s multiple comparisons test).

We next checked HSAT1 chromosomal locations by BLAST searching the GRCh38.p14 (GCA_000001405.29) and CHM13-T2T v2.0 (GCA_009914755.4) human genome assemblies. For this purpose, we chose one of the smaller clones as representative of the family (HSAT1 clone from herein). HSAT1 sequences mapped to chromosomes 3, 4, 8, 14, and 22 in the GRCh38.p14 assembly, while in CHM13-T2T v2.0 matching positions were present in chromosomes 3, 4, 8, and all acrocentric chromosomes (Supplemental Table S2). This result confirms the higher quality of the CHM13-T2T assembly, supporting the identification of HSAT1 arrays in the pericentromeric regions of chromosomes 13, 15, and 21, in agreement with previous reports (Kalitsis et al. 1993). Next, we queried the NCBI nucleotide collection with the HSAT1 clone sequence (Supplemental Table S3). In addition to the expected hits in CHM13-T2T v2.0 and previously described HSAT1 ptRI-6 clone (L01057.1), 61.2% of the hits belonged to unlocalized sequences from a sequencing project aiming to create a human population-specific genome assembly (Bioproject: PRJDB10452) (Supplemental Figure S1). This fact points to the recurrency of unplaced tandem repeats (like HSAT1) in attempts to assembly the human genome.

A multiple alignment between the pTRI-6 HSAT1 clone (L01057.1), the CHM13-T2T HSAT1A array (Lee et al. 2022; Nurk et al. 2022), and three of our cloned HSAT1 sequences (Figure 1A), representative of different sizes (~200, 580, and 880 bp) was performed using the Geneious software, revealing overall identities in the order of 80 % (Supplemental Table S4).

As satDNA genomic occurrence and persistence can be determined by long-range organization and structure (Vondrak et al. 2020), we further addressed HSAT1 periodicity from publicly available data. As mentioned, the analysis of the CHM13-T2T assembly with the NTRprism software tool, specifically developed for that purpose, reported that HSAT1 presents a higher-order organization with predominance of 9-mer NTRs (Nested Tandem Repeats) (Altemose et al. 2022). To expand on this information and assess the recurrency of HSAT1 organization in a reference/non-tumoral cell line used throughout this paper, we applied the same approach to nanopore long-read sequencing data from GM12878 cell line (Utah/CEPH pedigree) (Jain et al. 2018), The periodicity spectrum detected across a 497.1 Kb read is shown in Figure 1B. The most frequent periodicity identified corresponded to the HSAT1 42bp monomer, followed by the 378bp 9-mer array that was found to be the prevalent HSAT1 NTR in the chromosome 13 array of the T2T assembly (Altemose et al. 2022).

As a rule, tandem repeats are underrepresented in databases (Monlong et al. 2018; Ershova et al. 2020), leading to a gross undervaluation of their genome representativeness. The evaluation of copy number fluctuation between different genomes is particularly interesting as it can provide insights into the evolutive behavior of a given tandem sequence. Given the lack of information and wide-ranging molecular studies on HSAT1, we next evaluated the copy number of this sequence in sixteen human cell lines (tumoral and non-tumoral; Supplemental Table S5) from different tissues, by qPCR. Additionally, we assessed HSAT1 copy number in two human/murid somatic cell hybrids for human chromosomes 14 and 21. Monomer copy number showed considerable variability between cell lines, which seemed to be independent of tissue type and origin (tumor-derived or non-tumoral) (Figure 1C, Supplemental Table S6). For example, MDA-MB-231, MDA-MB-468, or MCF7 (cell lines derived from breast adenocarcinomas) all have a significantly higher copy number than MCF10A (non-tumorigenic), but the latter presents more copies than SK-BR-3 (breast adenocarcinoma). The same pattern of variation can be observed in the lung cancer cell line repertoire assessed in this study (H1299, A549, H1975, H2228, and PC9). Given the well-known connections between genome instability and human satDNAs (Balzano et al. 2021), it would be reasonable to expect HSAT1 copy number variation (CNV) to differ between normal and tumor cell lines. Nonetheless, the polymorphic nature of these sequences (greatly varying even between individual arrays) and their natural behavior as sources of CNV in genome evolution (Warburton et al. 2008; Black and Giunta 2018; Lower et al. 2018; Porokhovnik et al. 2021), limit any assessment of HSAT1 CNV contribution to instability in the absence of information about the starting point. In fact, satellite CNV is often so substantial that it is cytogenetically detectable between individuals (Miga 2019). In addition to the observed high HSAT1 copy number variation (CNV) in non-tumoral cell lines (MCF10A, GM12878, HDFn, and GM03417), the analysis of somatic cell hybrids (GM11535 and GM08854) also seems to underscore the relevance of populational polymorphism. Indeed, from early studies, HSAT1 was reported to be largely present in chromosome 13 and 21, and less represented in other acrocentric chromosomes (Kalitsis et al. 1993). However, we find that the GM11535 cell line, with a single copy of human chromosome 14 per cell, presents a higher HSAT1 copy number than GM08854, which has 1-5 copies of human chromosome 21/cell. The discrepancy between these results and past observations could be explained by significant inter-individual variation in the size of HSAT1 arrays. Excluding somatic cell hybrids, HSAT1 genome representativity in the analyzed human cell lines ranges between ~0.05 and 2.82%, with an average of 0.83% (Figure 1C, Supplemental Table S6). These results are in line with the estimation of 0.43% for HSAT1 sequences in the CHM13-T2T genome assembly. To further assess the reliability of our qPCR quantification, we compare our results for the GM12878 cell line with the available sequencing data using RepeatMasker (Smit et al. 2015). The estimated genomic abundance based on the sequencing dataset was of ~0.2% (Supplemental Table S7), within close range of our qPCR estimate of ~0.1% for the same cell line. Thus, it is fair to assume that the qPCR quantification method used in this work is able to estimate HSAT1 representativity with an accuracy comparable to the estimates obtained from sequencing data analysis.

Many lncRNAs, and more specifically satncRNAs, have been linked to tumor formation and progression, essentially by their abnormal level of transcription (Sanchez Calle et al. 2018; Gao et al. 2020). Thus, we next evaluated HSAT1 transcription in the same set of cell lines used for copy number assessment by RT-qPCR. Additionally, we quantified HSAT1 transcripts in three human tissues (mammary, small intestine, and brain). The collection of cell lines derived from breast adenocarcinomas (MCF7, MDA-MB-231, MDA-MB-468, and SK-BR-3) and the one derived from lung carcinoma (H1299, A549, PC9, H1975, and H2228) presented variable transcriptional profiles (Figure 1D, Supplemental Table S8). When comparing to MCF10A, some cancer cell lines (MDA-MB-231, MDA-MB-468, H1299, and A549) have aberrant overexpression of HSAT1, while others (SK-BR-3, PC9, and H2228) still overexpress HSAT1, but to a much smaller extent. This comparison is particularly relevant because of the same tumor type of the analyzed cell lines. Such varying levels of transcription in similar cancer tissues question the premise of a analogous overexpression behavior for HSAT1 ncRNA (Bartonicek et al. 2016; Chatterjee and Sengupta 2021), or at least presupposes a more complex regulatory scenario. This is especially true when comparing two cell lines derived from lung carcinomas, H1299 and H1975, the former expressing more than 1 million times more than MCF10A and the latter underexpressing HSAT1 (approximately half of MCF10A transcription). Other cancer cell lines – HeLa and HepG2 –show transcription upregulation, while Caco-2 has lower HSAT1 transcription. Non-tumoral cell lines – GM12878, GM03417, or HDFn – have more HSAT1 transcripts than MCF10A. Nevertheless, none of the analyzed non-tumoral cell lines show a transcription level comparable with the aberrantly expressing cancer cell lines (e.g., MDA-MB-468 or HeLa). By analyzing HSAT1 ncRNA in human tissue samples, we could observe a non-statistically significant difference between MCF10A (originated from mammary gland) and mammary tissue (as expectable), but a higher transcript level in small intestine and brain tissues.

### HSAT1 physical mapping

We next proceeded to physically map isolated HSAT1 clones in human chromosomes and compare the corresponding locations with available data. In Figure 2A we present FISH (Fluorescent *in situ* hybridization) results showing HSAT1 mapping in a metaphase preparation from GM12878. The tested subset of HSAT1 clones (representative of the three obtained sizes) showed very similar hybridization patterns in the same pericentromeric sites. Likewise, HSAT1 chromosomal locations were similar between cell lines (not shown). Clear hybridization signals (with different intensities) were observed in all acrocentric chromosomes. Moreover, we detected FISH signals in chromosomes 1 and 3 pericentromeric region, the former location (chr 1) reported here for the first time (as far as we know).

**Figure 2.**
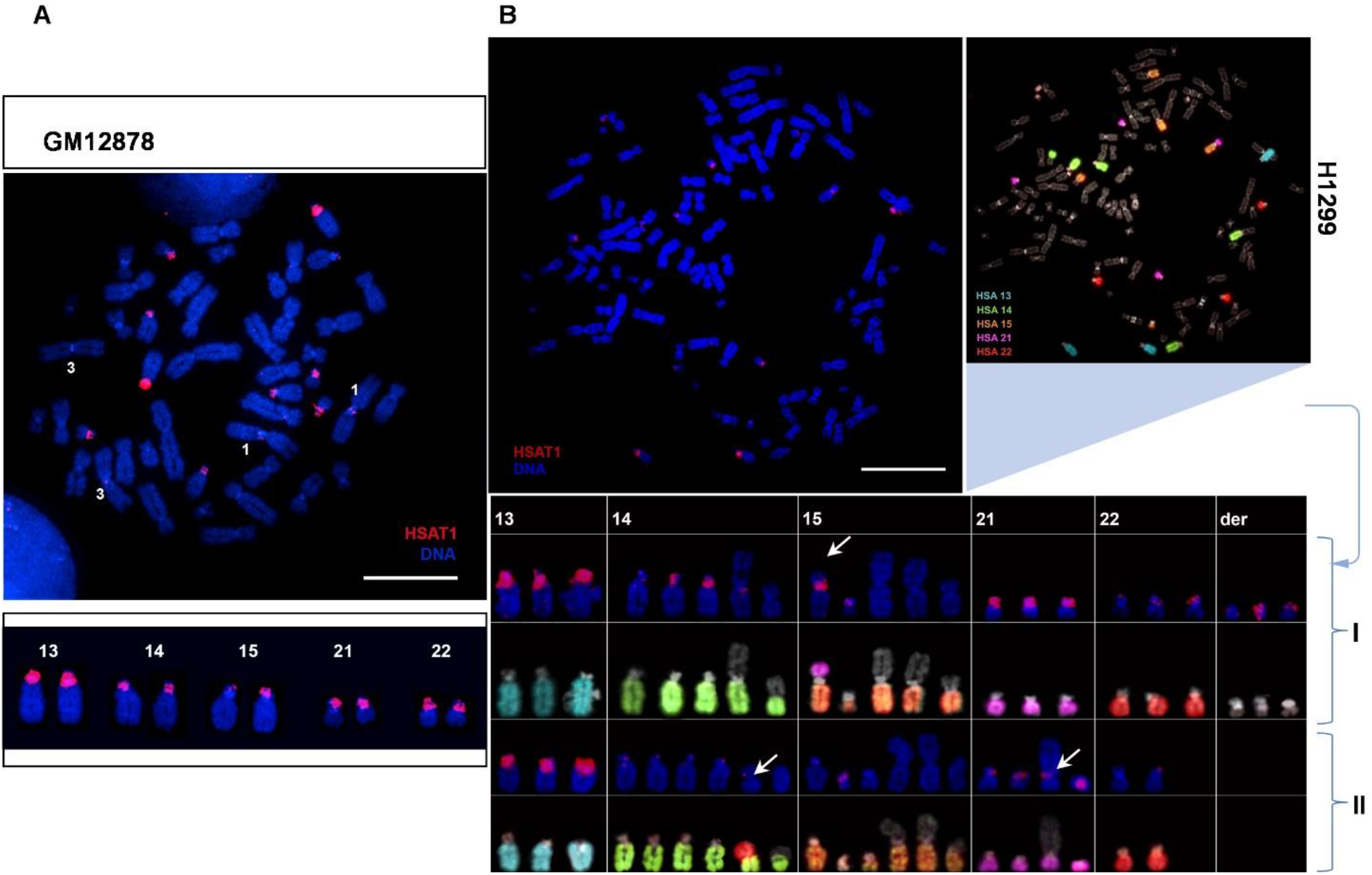
HSAT1 FISH mapping (red) in human chromosomes (blue). **(A)** – HSAT1 mapped in GM12878. Obtained hybridization signals in acrocentric chromosomes are highlighted above. Hybridization signals are also present in chromosomes 1 and 3. Chromosomes were identified by reverse-DAPI. **(B)** – HSAT1 mapped in chromosomes from H1299, sequentially hybridized with human painting probes for acrocentric chromosomes. Corresponding chromosomes are visible in the table above (two different clones). The illustrative metaphase corresponds to clone I (first two rows). The column “der” for clone I shows three derivative chromosomes (non-acrocentric) with visible HSAT1 signals. White arrows indicate chromosomal alterations occurring with acrocentric chromosomes and modifying HSAT1 hybridization pattern. Scale bars represent 10 μm.

We further mapped HSAT1 in the tumor cell line H1299, in which we detected the highest level of HSAT1 ncRNA transcripts. After HSAT1 probe hybridization, we performed sequential FISH with painting probes for human acrocentric chromosomes. Figure 2B illustrates two different clones found in this cell line, both with clear presence of chromosomal alterations (e.g., translocations), evidently affecting the intensity/presence of HSAT1 hybridization signals. Given these results, CNV between H1299 clones is also likely to be happening. The cytogenetic mapping analysis points to a possible role of HSAT1 as a player in genomic/chromosomal instability and a plausible fragile pericentromeric location, warranting further exploration.

### HSAT1 transcripts: Cellular Patterns

To further characterize HSAT1 transcription, we proceeded to determine the subcellular localization of HSAT1 transcripts by RNA-FISH, an approach that has been used to visualize and analyze the spatial distribution of ncRNAs in several species and conditions (Brown and Buckle 2010; Trofimova et al. 2014; Goenka et al. 2016; Ferreira et al. 2019a; Bury et al. 2020). After detecting HSAT1 ncRNA foci, we performed RNase A treatment prior to hybridization, to ensure signal specificity and exclude unintentional DNA hybridization (analysis shown in H1299 cells). RNA-FISH signals decreased significantly, as also seen by the evaluation of the average intensity of active fluorescent objects in control RNA hybridization and RNase treatment (Figure 3A; Supplemental Table S9). Detection of HSAT1 transcripts in different cell lines allowed to perform a single-cell topographic observation of the obtained signals (Figure 3B). RNA-FISH is presented in seven cell lines to show feasibly distinctive transcript features between cell lines with similar amounts of HSAT1 transcription. HSAT1 transcripts exhibit cluster-like organization and consistent nuclear localization, though with distinct signal patterns in different cell lines. For example, HeLa, H1299, and MDA-MB-231 cells (all aberrantly expressing HSAT1) have evident differences concerning the number and spatial distribution of HSAT1 ncRNA foci. Moving towards in our approach, we studied the nuclear localization of HSAT1 transcripts, coupling HSAT1 RNA-FISH with immunofluorescence (IF) for the detection of fibrillarin. Our intent was to compare nucleoli localization with HSAT1 ncRNA signal clusters. Image 3C illustrates this analysis in H1299 and MCF10A, where HSAT1 transcripts can be spotted contiguously to nucleoli (peripheric). Confocal 3D images allow to visualize signal spatial distribution (with isosurfaces and orthogonal slices for axis projections). We also performed sequential DNA-FISH to the slide with RNA-FISH, imaged the same slide fields and merged both confocal images (Figure 3D). HSAT1 signals are differently organized in both FISH experiments, even though some RNA signals colocalize with HSAT1 DNA, pointing to nascent HSAT1 transcription.

**Figure 3.**
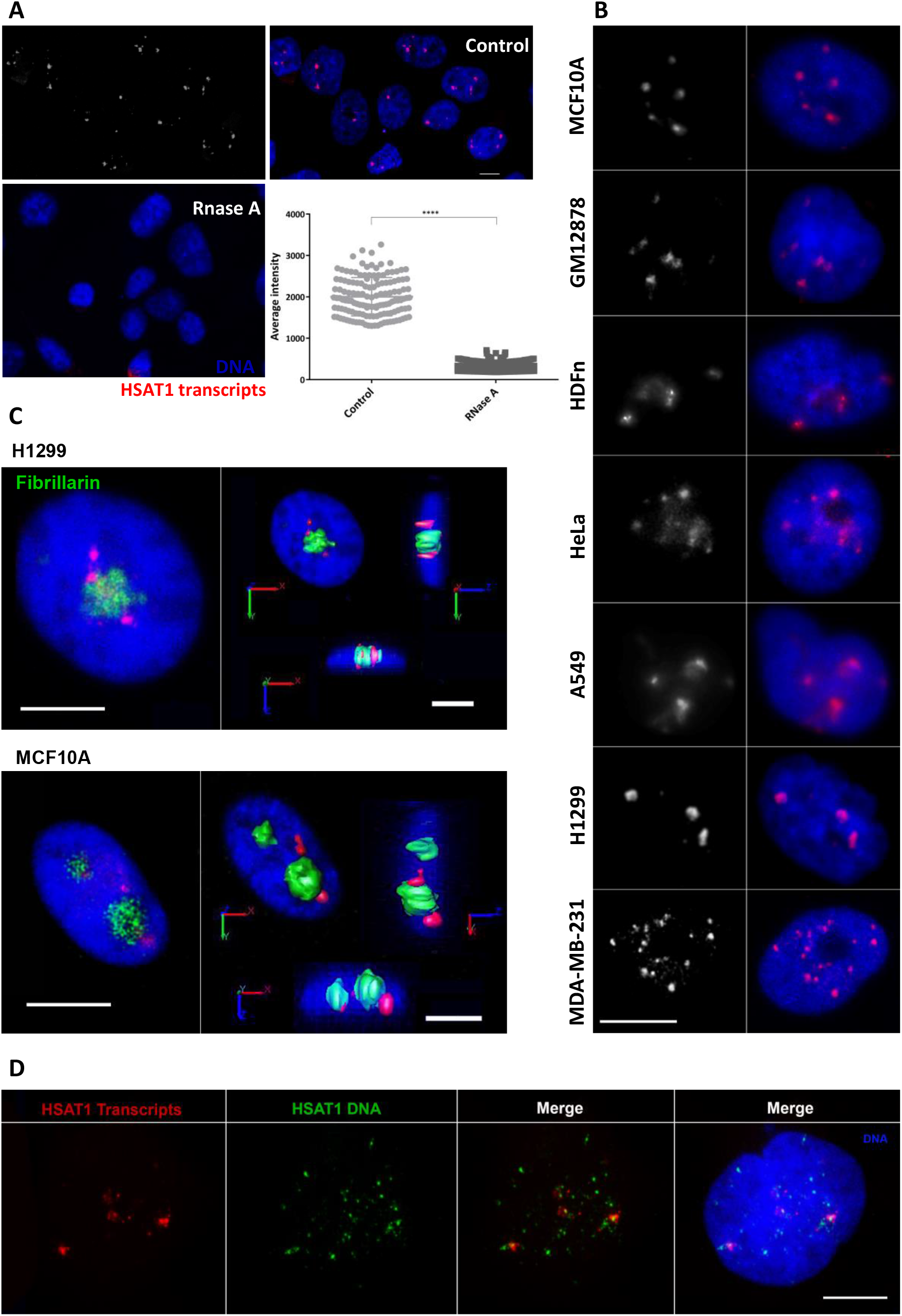
Detection of HSAT1 transcripts by RNA-FISH and RNA-FISH/IF. **(A)** – HSAT1 RNA-FISH with RNase A treatment. HSAT1 transcripts were detected by RNA-FISH (red) in control and treated cells. Signal decrease in RNase-treated cells demonstrates that the observed signals are RNA-specific. Evaluation of the average intensity of active signal objects (all slices) in RNA-FISH control and RNA-FISH + RNase A was performed in ‘Counting and Tracking’ (AutoQuant X3). Analysis shown in H1299. Values are mean ± SD. ****P≤0.0001 (Unpaired t test). **(B)** – HSAT1 nuclear organization of HSAT1 transcripts (red). Standard Streptavidin-Cy3 detection was used in highly expressing cell lines. Cell lines with reduced HSAT1 transcription required signal amplification (SuperBoost™ Tyramide Signal Amplification). Different spatial distribution and number of foci are observable between cell lines with similar amounts of HSAT1 transcripts (RT-qPCR data). **(C)** – Spatial organization of HSAT1 transcripts in relation to nucleoli. HSAT1 RNA-FISH (red) coupled with IF for fibrillarin detection (green). HSAT1 transcripts seem to accumulate adjacently to nucleoli, as seen by confocal 3D image analysis for H1299 and MCF10A cells. Orthogonal slices for axis projections are displayed with isosurfaces for both channels. **(D)** – HSAT1 RNA-FISH (red) followed by HSAT1 DNA-FISH (green). Merged confocal images show distinct signal features, with some co-localized signals. DNA is in blue (DAPI) in all the presented images. Scale bars represent 10 μm.

### Scanning the features of HSAT1 transcripts

Our approach to analyze HSAT1 transcripts followed by searching HSAT1 clones in NCBI SRA data, specifically data from PRJNA362590 (HeLa PacBio ncRNA-Seq; SRA experiment: SRX2505545). We obtained 307 hits for 128 target sequences, with a % identity superior to 80.8% and multiple reads bigger than 1 Kb. For verifying the possible existence of annotated lncRNAs with similarity with HSAT1, we next BLAST searched HSAT1 against LNCipedia Version 5.2 Full Database (Volders et al. 2019) (Supplemental Table S11). Our results retrieved three NONCODE v4 annotated transcripts with two exons: lnc-RNF170-2:1 (475 nt, sense intronic ncRNA), lnc-RNF170-1:1 (223 nt; lincRNA; NONCODE Gene ID: NONHSAG050120.2), and lnc-RNF170-5:1 (629 nt; lincRNA; NONCODE Gene ID: NONHSAG050119.2). By analyzing UCSC Genome Browser (Lee et al. 2022) (http://genome.ucsc.edu.) on human (GRCh38/hg38), annotated ncRNAs all map to chromosome 8, in overlapping locations with annotated SAR (RepeatMasker (Smit et al. 2015)) and three ESTs (Expressed Sequence Tags) from GENCODE v39 (Frankish et al. 2020). The latter sequences were found in nervous, liver, and testis tissue (Bu et al. 2015). The % of identities with HSAT1 clone were ~80%.

In order to have a deeper understanding of HSAT1 transcripts, we performed 3’ RACE (Rapid Amplification of cDNA Ends) (Yeku and Frohman 2011) in HeLa RNA using an oligo-dT anchor primer for reverse-transcription and a forward PCR primer targeting the HSAT1 sequence. Analysis of 3’RACE products by agarose gel electrophoresis revealed a ladder of products, with the most intense band around 550 bp (Figure 4A). This result suggests that HSAT1 transcripts are polyadenylated and thus, likely transcribed by RNA Polymerase II (Hirose and Manley 1998). To characterize the amplified products, we performed a 300nt paired-end high-throughput sequencing using the Illumina MiSeq platform. A total of ~1×10^5^ reads (Supplemental Table S11) were obtained (for R1 and R2), which were quality and size filtered and assembled into ~3.5 x10^4^ complete 3’-RACE transcripts, as described in Supplemental Methods. Approximately 70% of these assembled transcripts presented the HSAT1 motif and were distributed across a size range of 51 to ~400 nucleotides, with peaks corresponding to multiples of the 42-monomer size (Figure 4A). Although we cannot exclude PCR size-amplification bias, the most prevalent sequence size was ~170 nt. Within this universe, 16,332 sequences were found to be unique, attesting to the high complexity of the HSAT1 transcriptome. By analyzing sequence reads in a window of 200 nucleotides, we could observe a progressive reconstruction of longer reads by piercing together smaller ones (Figure 4B, Supplemental Table S12), possibly suggesting mechanisms of ACPA. To test this theory, we examined the structure of a representative read, having found multiple alternative and non-canonical PASs (Tian et al. 2005), organized in a known poly(A) signal structure (Proudfoot 2011), and cleavage sites often corresponding to the actual read lengths. To get a better perspective of the degree of sequence variability, we clustered this dataset into groups with a minimum sequence identity of 90%, identifying a total of 257 clusters, 50 of which had more than 50 elements (Figure 4D, 4E). We next unbiasedly searched for sequence motifs within each of the 50 mentioned clusters. We found that the obtained repeated motifs invariably compose, or were composed of, HSAT1 42 nt monomers (Supplemental Figure S3).

**Figure 4.**
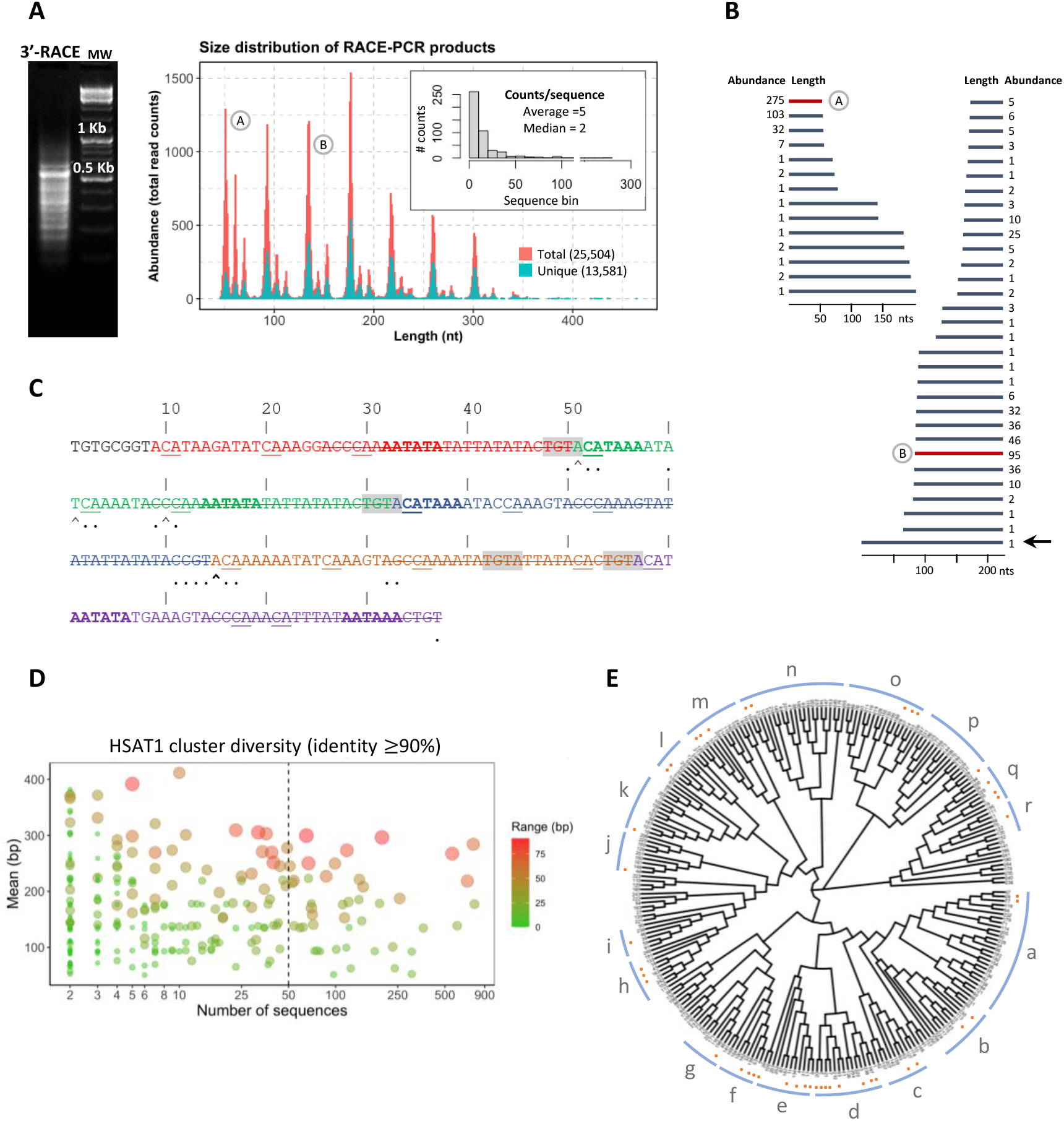
HSAT1 3’ RACE analysis. **(A)** - Agarose gel corresponding to HSAT1 3’RACE-amplified transcripts (left) (ladder of bands, smear-like); molecular weight (MW) (right). A size distribution plot is presented for the graphical representation of HSAT1 reads. Approximately 70% of the assembled transcripts contained the HSAT1 motif and were distributed across a size range of 85 to ~400 nucleotides, with peaks corresponding to multiples of the 42-monomer size. From the total of HSAT1 sequences, 16,332 sequences were found to be unique (Figure 4A, blue in size distribution plot). The bar chart in the top right corner addresses the high representation of unique sequences, visible in the distribution of counts/sequence. A and B (round) sequences are representative of the identified peaks and are displayed in B. **(B)** – Depiction of HSAT1 tandem transcript organization. Length and abundance are indicated. Unique sequences of smaller sizes are present within longer ones. In a universe of 200 nucleotides, it is possible to reconstruct transcripts of longer length. The black arrow points to the longer represented read (structure explored in C). **(C)** – HSAT1 read structure analyzed in the light of the consensus mammalian poly(A) signal. Different colors display HSAT1 monomers (42 nt). HSAT1 monomers are organized in alternative A (17 nt) and B (25 nt; strikethrough nucleotides in the figure) repeats. Sequences that may function as Poly(A) signals (PAS) hexamers (Tian et al. 2005) are highlighted in bold. The motif represented in shades of grey corresponds to the sequence that functions as the recognition of the poly(A) signal in the absence of the canonical hexamer element [A(A/U)UAAA]. Nucleotides located at the site of optimal 3’ cleavage (CA), named the poly(A) site, are underlined. Arrows point for the largest number of duplicates that are cleaved at that nucleotide position (bold for the largest most abundant). Dots represent the cleavage location of duplicates that contain a difference higher than 1 nucleotide from the previous sequence. The cleavage positions depicted in this representative read intend to emphasize the possible occurrence of Alternative Polyadenylation (APA), resulting in the observable variation of transcript length. **(D)** – HSAT1 transcript sequences were clustered, selecting a threshold of an identity score of 90% to determine the cluster membership. Clusters are organized by mean sequence size and number of sequences. The presented colors determine the range (bp) between sequences of the same cluster. **(E)** – Phylogenetic tree depicting transcript variability, constructed from the multiple alignment between the center sequences of each cluster. Clusters can be grouped accordingly to their distance (groups a-r). Orange dots represent clusters with more than 50 elements, selected for posterior motif discovery.

## Discussion

From old parallelisms with ‘darker’, ‘unknown’, or ‘useless’ contexts, satDNA has been growing amongst research interests, especially when related analysis obstacles are progressively being outpaced. Centromeric and pericentromeric satellite sequences were effectively difficult to incorporate in genomic assemblies, as linearly order long arrays of tandem repeats was technically unachievable (Rudd and Willard 2004; Miga 2021; Miga and Alexandrov 2021). This situation is observable in the initial part of this paper: isolating and subsequently mapping HSAT1 clones in the human reference genome GRCh38.p14 highlighted the still significant assembly gaps in reference acrocentric/pericentromeric regions. When comparing reference mapping with our FISH results in GM12878 (Figure 2A), we can clearly see that HSAT1 should be highly represented in *in silico* maps of chromosomes 13 and 21 (intense FISH signals but absent in reference sequences). Indeed, the entire p-arms of acrocentric chromosomes (and therefore, HSAT1 repeats) represented long stretches of gaps, until the emergence of CHM13-T2T (Antonarakis 2022). The difficulties in assembling HSAT1 repeats (AT-rich and highly shared between acrocentric chromosomes) were experienced while sequencing acrocentric p-arms (Nurk et al. 2022) and are observable in our BLAST search returning 61.2% of unlocalized sequences (PRJDB10452). CHM13-T2T was an unquestionable addition to satDNA studies - our HSAT1 clone mapping in CHM13-T2T (Supplemental Table S2) was consonant with our FISH mapping in acrocentric chromosomes (Figure 2). FISH signals were also observable in chromosomes 1 and 3. In particular, HSAT1 mapping in chromosome 1 (not previously reported) points to the need of inquiring human satDNA variation. Also, it suggests the pertinence of using cytogenetic mapping in both clinical and non-clinical contexts.

SatDNA sequences can represent up to 10% of the human genome (Podgornaya et al. 2018; Puppo et al. 2020) and have a major representation of CNVs, even between individuals (Warburton et al. 2008; Black and Giunta 2018; Lower et al. 2018). These changes in satellite copy number may occur through arrays contractions/expansions, due to recombination between repeats with unequal change (Plohl et al. 2008; Lower et al. 2018). HSAT1 monomer copy number quantification in a set of human cell lines likely features this polymorphism as a source of human variation. For example, αSAT representativity can vary between 1-5% and HSAT2/3 between 1-7% (different human populations) (Altemose et al. 2014; Miga 2019). So, the observed differences in HSAT1 copy number between cell lines (of 2.5% in some cases) was quite expectable and might not be related with the characteristics of each cell line and/or the use of cancer cell lines. Satellite CNV in cancer can be particularly advantageous for polymorphic variation between cell populations and rapid evolution (de Lima et al. 2021). However, human genome variation might be in play here, which calls for the need of assessing satDNA CNV in a pangenomic approach (Logsdon et al. 2021; Wang et al. 2022), in order to evaluate its functional significance in diverse states, evidently assuming that we might still being underestimating the genomic representation of these sequences.

By turn, (peri)centromeric transcription has been related with cancer (Ting et al. 2011; Sanchez Calle et al. 2018; Bury et al. 2020; Nogalski and Shenk 2020; Chatterjee and Sengupta 2021). HSAT1 transcription was assessed in the same cell lines used for copy number analysis, yet with even more dissimilar results. So, in concordance with the possible polymorphic nature of this sequence, there was no clear evidence of an association between HSAT1 copy number and expression, as already seen in other works with satellite sequences (Ferreira et al. 2019b). When quantifying HSAT1 transcripts, we analyzed cancer cell lines from similar origins with astounding different profiles. In cancer, ncRNAs and tumor suppressor genes/oncogenes share the trait of highly variable transcription, which can indicate the presence of post-transcriptional complexed-regulated pathways (Carter et al. 2021). Furthermore, the tendency for mutation of genomic regulators in diseases such as cancer could have a determining role in ncRNA transcription deregulation (Zhou et al. 2016; Pratt and Weng 2018; Carter et al. 2021). Thus, the variable genetic and epigenetic landscape of HSAT1 might be determining transcription differences. For example, changes in histone methylation, particularly in the levels of H3K9me3 (a typical repressive mark found in satellite heterochromatic regions) (Déjardin 2015; Vojvoda Zeljko et al. 2021), can relate with cancer predisposition (Peters et al. 2001; Nakagawa and Okita 2019). If HSAT1 transcription turns to be a player in genome instability and epigenetic regulation, more functional studies are needed in order to place the transcriptional event - causing changes in epigenetic regulators (Hall et al. 2017) or being deregulated as a consequence. This duality can also be related with cancer chromosomal instability, as shown in Figure 2B. Chromosome breaks are more prone to occur in satellite pericentromeric regions (Black and Giunta 2018), causing chromosomal rearrangements with the ability to compromise genome stability (Balzano et al. 2021) and altering the transcription status of genes or even satellite sequences themselves (Fournier et al. 2010; Ferreira et al. 2015; Puppo et al. 2020).

Through RNA-FISH experiments in multiple cell lines, we subsequently address the subcellular localization of these transcripts and found that they are nucleus-specific, despite having different signal topographies in different cell lines. It became clear that HSAT1 transcription deregulation does not result in altering transcript location, although possibly resulting in a different organization within the nucleus. HSAT1 RNA-FISH together with fibrillarin staining allowed to address a perinucleolar presence to HSAT1 ncRNA foci. The signal distribution of HSAT1 can be linked with the organization of satellite DNA and RNA into chromocenters, often associating in close proximity to the nucleoli and/or nuclear membrane (Camacho et al. 2017; Brändle et al. 2022). Still, epigenetic alterations and/or changes in nuclear architecture could explain different patterns, like the one seen in HSAT1 transcripts from MDA-MB-231 (Figure 3C).

3’RACE allowed us to perform a deeper characterization of HSAT1 transcripts by proving the accumulation of polyadenylated ncRNAs. Other reports show that both polyadenylated and non-polyadenylated pericentromeric transcripts can be detected in human (Dobrynin et al. 2020). Polyadenylation of Pol II ncRNA transcripts can be tightly related with transcription termination, and intensively regulated in the presence of cellular stresses and/or cancer-associated mutations (Proudfoot 2016; Tian and Manley 2017). The size variability of HSAT1 transcripts, also attested by the presence of multiple PASs, is likely to be a result of APA. APA can be possibly regulated by the levels of U1 telescripting (when high, inhibiting PCPA; when low, resulting in smaller transcripts) (Cugusi et al. 2022). Moreover, poly(A) site selection becomes more complex in the presence of non-canonical hexamers (Beaudoing et al. 2000). By exploring our transcript variability, we can probably assume the absence of preferable loci for the transcription of these pericentromeric sequences. In any case, accumulation of HSAT1 transcripts presumably depends on the joined complexity of transcriptional and post-transcriptional pathways (Eymery et al. 2009b).

In the future, the association of HSAT1 with the plethora of functional significant satncRNAs would benefit from coupling our analysis with 5’RACE or sequencing of Pol II transcription start sites (TSSs), in order to obtain deeper information regarding transcript full size and splicing complexity (Lagarde et al. 2016; Yan et al. 2022).

Irrespective of being a consequence of chromosomal rearrangements and/or changes in (epi)genetic cell states, the abnormal transcription of α satellite and classical satellite sequences is a trait of tumor cells (Eymery et al. 2009a; Hall et al. 2017; Puppo et al. 2020). The present work highlights the former statement, trailing the way for the characterization of HSAT1 transcripts. More functional work is crucial, possibly initiating by HSAT1 knockdown and posterior evaluation of cellular phenotypes (Ramilowski et al. 2020): determining if this satellite is, for example, possibly involved in organizing genome architecture (as several tandem repeats) (Muller et al. 2019; Balzano et al. 2021), or in gene expression regulation during stress, development, and pathology (Feliciello et al. 2021). Studying the upstream regulation of HSAT1 transcripts is also essential for the mechanistic understanding of involved cellular pathways (Gao et al. 2020). This paper closes a lacuna in HSAT transcription studies, since it proves that, like the other classical satellites (HSAT2/3), HSAT1 is indeed transcribed and most likely intensely deregulated depending on the gambling of cancer (epi)genetic cause-effect scenarios.

## Material and Methods

### Cell culture, chromosome harvesting, genomic DNA/RNA isolation

Cell culture specificities are described in Supplemental Methods. Chromosome harvesting procedures were routinely followed. Genomic DNA extraction was achieved with QuickGene DNA Tissue Kit S (Fujifilm Life Science) (instructions accordingly). RNA was isolated following the mirVana Isolation Kit (Ambion, Thermo Fisher Scientific). Total RNA was purified from DNA using the TURBO DNA-free ™ Kit (Ambion, Thermo Fisher Scientific). DNA and RNA were quantified with Qubit™ dsDNA BR Assay Kit and Qubit™ RNA BR Assay Kit, respectively. Total RNA pools from human brain (cat. no. 636530, Takara Bio USA, Inc), small intestine (cat. no. 636539, Takara Bio USA, Inc), and mammary gland (cat. no. 636576, Takara Bio USA, Inc) were also used for the assessment of HSAT1 expression in different tissues.

### HSAT1 isolation, cloning, and *in silico* analysis

HSAT1 was amplified from human genomic DNA with two sets of specific designed primers. Primers were designed using Primer 3 (Rozen and Skaletsky 2000), available in Geneious R9 version 9.1.8 (Biomatters); and are described in Supplemental Table S1. PCR program was as following: initial denaturing step at 94°C for 10 min; 30 cycles of 94°C for 1 min (denaturation), 57°C for 45 s (annealing) and 72°C for 45 s (extension); final extension at 72°C for 10 min. PCR products were run in an agarose gel and the obtained bands were isolated and cloned. Specific steps for cloning and clone *in silico* analysis/mapping are detailed in Supplemental Methods.

For the quantification and periodicity study of HSAT1, sequencing data was extracted from the Whole Human Genome Sequencing Project, NA12878 (https://github.com/nanopore-wgs-consortium/NA12878). The header of each read was renamed by numerical order (from 1 to 15666888). The HSAT1 repetitive sequence (SAR, accession DF0001062.4) was extracted from the Dfam database (https://dfam.org/). HSAT1 clone repetitive sequence was assembled in-house. Quantification was performed using RepeatMasker (Smit et al. 2015) to detect the presence of HSAT1 (SAR) and of HSAT1 clone on reads from NA12878 sequencing data. Genomic abundance for each of the repetitive sequences was estimated based on number of masked bases / total bases from reads * 100. For the periodicity studies of HSAT1 repetitive sequences, reads with a number higher than 400 Kb containing only these sequences were selected. Then, the NTRprism (https://github.com/altemose/NTRprism) scripts were used on the selected reads.

LNCipedia annotated lncRNAs from Version 5.2 Full Database (Volders et al. 2019) were download (https://lncipedia.org/) and used as a custom BLAST database in Geneious.

### Metaphase Fluorescent *in situ* hybridization (Metaphase FISH)

In order to physically map HSAT1, FISH was performed as described in (Gonçalves et al. 2019) with slight modifications. Metaphase slides were treated with acetone for 10 min, baked at 65°C for 1 hour and denatured in an alkaline denaturation solution (0.5 M NaOH, 1.5 M NaCl) for 1-4 min. Clone probes were PCR labelled with biotin-16-dUTP (from Roche Applied Science). Hybridization was performed over-night. Post-hybridization stringency was done with 1×SSC at 73°C. Biotin-labelled HSAT1 probe was detected with Streptavidin Cy3 conjugated (Sigma-Aldrich). Preparations were mounted using Vectashield containing 4’-6-diamidino-2-phenylindole (DAPI) (Vector Laboratories) to counterstain chromosomes.

FISH with human acrocentric chromosome paint probes was performed sequentially to the hybridization with HSAT1 probe. Briefly, FISH slides were washed in 2×SSC for 10 min, followed by denaturation in alkaline denaturation solution (0.5 M NaOH, 1.5 M NaCl) for 2-5 min. The paint probes were labeled by DOP-PCR as follow: human 13 was labelled with Atto-488-dUTPs, human 14 with Atto425-dUTPs, human 21 with Atto-Cy5xx-dUTPs, human 22 with Atto-594-dUTPs and human 15 was labelled with Atto-425-dUTPs and Atto-594-dUTPs (Jena Bioscience). Hybridization was carried out overnight at 37°C. Post-hybridization washes were done as described above.

FISH images were observed using a Zeiss ImagerZ2 microscope coupled to an ORCA-Flash 4.0 digital camera (Hamamatsu) and captured using SmartCapture 4 software (Digital Scientific, UK). Digitized photos were prepared for printing in Adobe Photoshop (version 7.0).

### RNA-FISH/ RNA-FISH-IF/Sequential DNA-FISH

RNA-FISH was performed accordingly to (Ferreira et al. 2019a), with some modifications. Cells were hybridized overnight at 37°C with the PCR-Biotin-labelled HSAT1 probe. Probe detection was carried out with Streptavidin, Cy3 conjugated, or using Invitrogen™ Alexa Fluor™ 555 Tyramide SuperBoost™ Kit, streptavidin (Thermo Fisher Scientific), according to provided instructions. The second approach was used in cell lines with smaller amounts of HSAT1 transcripts (not visible with standard detection methods). When RNA-FISH experiments were coupled with immunofluorescence (IF) with anti-fibrillarin antibody, primary incubation was performed for 1h (anti-fibrillarin monoclonal mouse, 1:100, MA3-16771, Thermo Scientific), followed by incubation with secondary antibody (anti-mouse monoclonal FITC, 1:200, 81-6511, Zymed). Cells were then mounted with coverslips and counterstained with Vectashiled mounting medium containing 4’-6-diamidino-2-phenylindole (DAPI) (Vector Laboratories). RNase A treatments (0.8 mg/mL, Sigma-Aldrich) were performed after permeabilization for 3h at 37°C. Analysis of the average intensity of active objects was performed in 20 cells from control RNA hybridization and RNase treatment in AutoQuant X3 software (Media Cybernetics), ‘Counting and Tracking’ tool in all slices. For the sequential DNA-FISH, RNA-FISH slides were washed in 2×SSC for 10 min, treated with RNAse A (100 μg/ml in 2×SSC) for 30 min at 37°C, dehydrated through an ethanol series, denatured in 70% Formamide /2×SSC for 2 min at 72°C, and hybridized overnight with HSAT1 probe labelled with Atto488-dUTPs (Jena Bioscience). Post-hybridization stringent washes were done with 0.1×SSC at 42°C. Slides were mounted with Vectashiled mounting medium containing 4’-6-diamidino-2-phenylindole (DAPI) (Vector Laboratories).

For RNA-FISH/RNA-FISH-IF/sequential DNA-FISH images, confocal fluorescence imaging was performed on an LSM 510 META with a Zeiss Axio Imager Z1 microscope and LSM 510 software (version 4.0 SP2). Applied settings were constant. Used lasers were argon (488 nm) set at 12.9%, helium-neon (543 nm) set at 50.8% and diode (405 nm) set at 9.9%. The pinhole was set to 96 mm (1.02 airy units) for argon laser, 102 mm (0.98 airy units) for helium–neon laser, and 112 mm for the diode laser using a 63 × objective. Images were captured at a scan speed of 5 with 1 μm thick Z sections. AutoQuant X3 software (Media Cybernetics) allowed 3D deconvolution. Subsequent TIFF processing was run in ImageJ (1.52v). The three-dimensional isosurfaces and orthogonal slices (perpendicular or parallel angles) were produced in Image Pro Premier 3D (version 10, Media Cybernetics).

### DNA copy number absolute quantification (quantitative Polymerase Chain Reaction-qPCR)/ RNA expression analysis by Real-Time Reverse Transcriptase −qPCR (RT-qPCR)

HSAT1 copy number evaluation was performed using a standard curve obtained with serial dilutions of the recombinant plasmid (with HSAT1 clone). This method allowed to interpolate CT values (obtained in different DNA samples) against the standard curve. Real-time qPCR reactions were performed with MeltDoctor HRM Master Mix (Applied Biosystems, Thermo Fisher Scientific), according to manufacturer’s protocol. The standard curve method was also used for HSAT1 ncRNA quantification, as previously validated (Chaves et al. 2017). Quantification was carried using SensiFAST™ SYBR® Hi-ROX One-Step Kit (Bioline). Absolute RNA quantification was obtained by interpolating its CT value against the standard curve. All reactions used 100 ng of RNA and all data were normalized to MCF10A.

Primers, standard curve parameters, and PCR programs for both experiments are present in supplemental material (Supplemental Table S13). Primer specificity was always evaluated by the generation of a melt curve. All reactions were performed in triplicate, and negative controls were also included in the plate. Data were analyzed using the same parameters on the StepOne software (version 2.2.2, Applied Biosystems, Thermo Fisher Scientific). All data are presented as mean ± standard deviation. Statistical significance was determined using ANOVA tests. ns (non-statistically significant) for p > 0.05, *p ≤ 0.05, **p ≤ 0.01, ***p ≤ 0.001, ****p ≤ 0.0001.

### 3’ RACE, RACE-Seq, and sequencing analysis

Further characterization of HSAT1 transcripts (size and poly-A tail) the 5’/3’ RACE Kit, 2nd Generation (Roche) was followed. HeLa cDNA was prepared using the kit and subjected to PCR using the provided PCR anchor primer and the HSAT1 forward primer (Supplemental Table S1, F1). 3’RACE was coupled with high-throughput sequencing, performed by STAB VIDA NGS sequencing service. The analysis of the generated sequence raw data was carried out using CLC Genomics Workbench 12.0.3 (https://www.qiagenbioinformatics.com/). The used protocols for sequence analysis are detailed in Supplemental Methods. The scripts and produced data are publicly available in https://github.com/GamaPintoLab/HSAT1A-transcript-analysis.

## Data Access

Sequence data for HSAT1 clones is available in GenBank under accession numbers: OP172545 - OP172627. The 3’ RACE-Seq data generated in this study have been submitted to the NCBI BioProject database (https://www.ncbi.nlm.nih.gov/bioproject/) under accession number PRJNA867346.

## Competing Interest Statement

The authors declare no competing interests.

## Acknowledgments

This work was supported by EXPL/BIA-OUT/1028/2021 project grant from FCT, Portugal (to S.L. and R.C.), the Scientific Employment Stimulus contract CEECIND/01825/2017 (to S.L.), and UIDB/04046/2020 and UIDP/04046/2020 Centre grants (to BioISI), all from FCT, Portugal. M.L. is recipient of a PhD fellowship (Ref SFRH/BD/147488/2019) from FCT (Portugal).

## Author Contributions

M.L., S.L., and R.C. conceived and designed the original idea. M.L. wrote the first manuscript draft, conducted the experiments, analysis, and results preparation. D.F. collaborated with experiment design. S.L. assisted closely in experiment performance and performed some of the experiments. G.V. collaborated in some experiments. D.E. performed HSAT1 representativity and NTR analysis. M.G-C. and D.E. performed RACE-seq analysis. M.L., S.L., M.G-C. and R.C. wrote the final manuscript.

## Notes

### Competing Interest Statement

The authors have declared no competing interest.

